# Long-term patterns of ultra-low sulfur fuel oil bioremediation in Arctic shorelines using *in situ* mesocosms

**DOI:** 10.64898/2026.08.27.747307

**Authors:** Esteban Góngora, Ya-Jou Chen, Nastasia J. Freyria, Antoine-O. Lirette, Charles W. Greer, Lyle G. Whyte

**Affiliations:** Natural Resource Sciences, McGill University, Sainte-Anne-de-Bellevue, Quebec, Canada; Division of Natural and Applied Sciences, Duke Kunshan University, Jiangsu, China; Energy, Mining and Environment Research Centre, National Research Council Canada, Montreal, Quebec, Canada

**Keywords:** Ultra-low-sulfur fuel oil, biodegradation, beach sediment, Northwest Passage, microbiome

## Abstract

New maritime regulations restricting high-sulfur fuels have led to the transition to new low sulfur fuel oils (LFSOs). We do not know how LSFOs will behave in marine environments and how they will respond to available remediation strategies, presenting an environmental risk. The risk will be even higher in the remote high Arctic, especially along the Northwest Passage (NWP), for which an increase in shipping traffic is expected by the end of the century. In this study, we evaluated the long-term (one year) biodegradation potential of the native microbial community of NWP beach sediment using *in situ* mesocosm experiments with two different types of LSFOs: a marine gas oil (Marine diesel) and an ultra-low sulfur fuel oil (ULSFO). We observed that the lighter Marine diesel was biodegraded better (72.0%) than the heavier ULSFO (32.5%). We described composition of the microbial community of the mesocosms using 16S rRNA gene amplicon sequencing and observed a decrease in microbial diversity for the fuel-treated samples compared to the untreated controls. Despite the decrease in overall diversity, we observed significantly higher abundances of known hydrocarbon degrading microbes (e.g., *Oleispira*, *Altererythrobacter*, *Gilvibacter*, *Pseudohongiella*) in the fuel mesocosms. Our study showed the potential to implement biodegradation as a remediation strategy under the cold and oligotrophic environmental conditions present throughout the NWP. However, we also observed that microbes on their own cannot degrade the entirety of the removed fuel and other types of remediation will need to be considered to complement the natural biodegradation observed here.

**Highlights:**

- The natural attenuation of low sulfur fuels under Arctic conditions was characterized
- Marine diesel was more easily biodegraded than ULSFO
- Arctic beach bacteria can adapt to the presence of oils on the shorelines
- These microbes cannot fully biodegrade the tested fuels under Arctic conditions

## Introduction

In 2020, new regulations from the International Maritime Organization (IMO) came into force which set a maximum global cap for sulfur content in ship fuels of 0.5% for open seas and 0.1% within sulfur emission control areas (SECAs) located close to highly populated zones (Vedachalam et al., 2022). These changes have caused the shipping industry to phase out from the use of heavy fuel oils (HFOs; < 3.5% sulfur content) into global cap- and SECA-compliant fuels which include marine diesels and a new generation of low sulfur fuel oils (LSFOs) which can be further categorized as very low sulfur fuel oils (VLSFOs; < 0.5% sulfur content) and ultra low sulfur fuel oils (ULSFOs; < 0.1 % sulfur content). LSFOs comprise a wide variety of products with different hydrocarbon compositions (Daling and Sørheim, 2020; Faksness et al., 2024; Nelson et al., 2022; Yang et al., 2023). These differences are caused by the processes used by refineries to produce LSFOs which include catalytic and thermal cracking, hydrodesulfurization, or blending of heavy fuel oils with low sulfur hydrocarbons (Faksness et al., 2024; Faragher et al., 2024; Kass et al., 2022; Vedachalam et al., 2022; Zou and Yang, 2023).

Climate change is causing a rise in temperatures globally, but especially in the Arctic which leads to a reduction in sea ice coverage and the possibility of summers with ice-free Arctic waters by the middle of the century (Shen et al., 2023). This presents an enticing opportunity for the shipping industry as ice-free conditions will lead to the opening of the Northwest Passage (NWP), a shorter navigation route through the Canadian Arctic Archipelago (CAA) which connects the northern Atlantic and Pacific Oceans (Howell et al., 2023). The ambiguity on the applicability of Canadian and international legislation to regulate shipping through the NWP (Bartenstein, 2019) along with an expected increase of drifting ice coming from the highest regions of the CAA into the NWP channels (Howell et al., 2023; Mudryk et al., 2021) and the rise of storms and waves in the region (Liu et al., 2016) will pose unsafe conditions for vessels going through the NWP. Additionally, the IMO approved a proposal to create a new SECA in the Canadian Arctic in April 2024 (International Maritime Organization, 2024). This is of importance as ships going through the NWP could be required to use ULSFO which leads to a possibility of these new type of poorly understood fuels to be released into the environment in the case of a spill.

Cleanup strategies such as skimmers, *in situ* burning, dispersants, or sorbent booms may not be effective for LSFOs (Daling and Sørheim, 2020; IMAROS, 2022). Additionally, it has been observed that spilt VLSFO can be neutrally buoyant and moves right under the surface which could reduce its detection capacity (Pålsson et al., 2024) and lead to a significant portion of the fuel reaching the shoreline. The geographical isolation of the highly uninhabited NWP will also slow and limit spill response in Arctic marine environments (Emmerson and Lahn, 2012). Because of these reasons, simpler remediation solutions, such as bioremediation, should be considered. Previous studies have documented the presence and hydrocarbon biodegradative capacity of Arctic microorganisms from marine environments such as seawater (Brakstad et al., 2015; Cao et al., 2022; Gofstein and Leigh, 2023; Gomes et al., 2022; Kampouris et al., 2023; McFarlin et al., 2014; Pyke et al., 2023; Vergeynst et al., 2019b), sea ice (Garneau et al., 2016; Lofthus et al., 2021; Vergeynst et al., 2019a), and beach sediments (Bragg et al., 1994; Y.-J. Chen et al., 2024; Durand et al., 2023; Ellis et al., 2022; Freyria et al., 2024; Garrett et al., 2003; Góngora et al., 2024b; Lirette et al., 2024; Prince et al., 2003). These studies show the feasibility of bioremediation as a primary cleanup strategy.

Nevertheless, the inhospitable, oligotrophic, and cold environment of the high Arctic hinders microbial activity (Durand et al., 2023; Gomes et al., 2022; Hunnie et al., 2023; Schreiber et al., 2023). A previous study from our research group (Góngora et al., 2024a) performed an *in situ* mesocosm study to test the biodegradation capabilities of the microbial community of NWP beach sediment treated with marine diesel, ULSFO, or HFO. After 1 month, higher biodegradation activity was observed in the low sulfur fuels, but particularly for ULSFO, only 47.6% of the fuel was removed from the mesocosms. Here we present a follow-up study to Góngora et al. (2024a) with the objective to test whether prolonged incubation times and the addition of nutrient amendments are able to improve fuel biodegradation. We hypothesized that extending the duration of the experiment to a year would help to account for the slow metabolic rates of microbes inhabiting near 0 °C and often below. We observed a very defined shift in the 16S rRNA gene community composition of the fuel treatments, which indicated that the microbiome was able to adapt to the presence of fuel. This was accompanied by an increase in the percentage of ULSFO removed after a year (62% of the fuel removed), out of which only 32.5% of the fuel was removed by biodegradation. Additionally, we hypothesized that the addition of nutrient amendments in the form of N and P fertilizers would stimulate the microbial community by replenishing the naturally low nutrient concentrations we previously observed in various NWP beaches (Góngora et al., 2024b). However, we did not observe a significant effect of the nutrient amendments on biodegradation or removal effectiveness. These results show that the Arctic environments across the NWP could be under imminent environmental threat if a spill were to occur and that it could take years for a shoreline to recover on its own.

## Materials and methods

### Sampling site description and experimental design

Assistance Bay (74.6509° N, 94.2983° W) is an uninhabited beach located approximately 17 km south-east from the hamlet of Resolute on Cornwallis Island, Nunavut, Canada. We chose this location for our experiment because Resolute is expected to be a key stopover hub in the NWP as the route becomes more widely used once sea ice coverage is reduced. Additionally, Assistance Bay faces the NWP so it represents one of the many remote beaches that could be impacted by a fuel spill from a ship travelling through the NWP.

The mesocosms were prepared as described by (Góngora et al., 2024a) with minor modifications. Fluortex netting (product reference 09-250/39, Sefar) pieces (15 × 15 cm) were washed with HPLC grade dichloromethane (DCM; Applied Biosystems) and attached to 15 × 15 × 0.5 cm stainless steel plates using nylon fishing line. The mounted netting was then saturated with Marine diesel (Glencore Limited) or ultra-low sulfur fuel oil (ULSFO; Shell Trading Rotterdam B.V.). A biostimulation treatment was also added in the form of a combination of an inorganic fertilizer (monoammonium phosphate; MAP; Sigma-Aldrich) and an oleophilic fertilizer (S-200 OilGone; S200 hereafter; International Environmental Products). A combination of these two fertilizers was chosen based on a previous study from our research group which found that both fertilizers led to a higher hydrocarbon removal compared to either of them applied individually (Y.-J. Chen et al., 2024). MAP was added at a concentration of 0.25 g kg^-1^ (Bell et al., 2013) and assuming that approximately 1 kg of sediment surrounds the netting (0.25 g per mesocosm) and S200 was added at a 1:1 ratio to hydrocarbon (following the manufacturer’s instruction) and assuming that approximately 3 g of fuel were added to the netting (3 ml per mesocosm). A control only using fertilizers but no added fuel was also included along with an environmental control with no fuel or fertilizer applied. Unlike the one-month experiment where the mesocosms were placed on a sheltered part of the beach, for this study we selected a section of the beach that is fully exposed to tidal action. The treatments and controls were deployed in duplicates on August 7, 2021 on two parts of the tidal zone: the supratidal zone which does not experience frequent tidal activity and the intertidal zone which is regularly submerged during the high tide. Each mesocosm was buried at a depth of 5 cm and with a distance of approximately 25 cm between each plate and replicate sets were separated by approximately 1 m. A time 0 (T_0_) control consisting of a mesocosm with added fuel was buried as described above, then immediately removed, and processed as described below.

Samples were recovered after a year (387 days) on August 28, 2022. Each mesocosm was individually packed in a sterile Whirl-Pak bag and taken to the Polar Continental Shelf Project laboratory in Resolute where they were immediately processed. Under aseptic conditions, the netting was separated from the metal plates and cut into four 7.5 × 7.5 cm pieces. Two of the fragments were individually rolled and placed inside 20 ml amber glass vials with closed caps with silicone liners (Thermo Scientific) and stored at −20 °C. The other two fragments were individually rolled and inserted into sterile 15 ml Falcon tubes. DNA/RNA Shield (Zymo Research) was added into the tubes until the netting was fully submerged and the tubes were stored at −80 °C. The tubes and vials were transported in coolers to McGill University (Montréal, Canada) where the amber vials were stored at −20 °C until they were processed for hydrocarbon analysis and the Falcon tubes were stored at −80 °C until processed for DNA/RNA extractions.

### Hydrocarbon analyses

Samples were sent to the Bigelow Laboratory for Ocean Sciences for hydrocarbon quantification of the mesocosms. Samples were analyzed by GC/MS using a modified EPA method 8270D as described elsewhere (Aeppli et al., 2018). A time zero (T_0_) control for all fuel and tidal zone combinations prepared the same way as the treatment mesocosms (as described above) was also sent for analysis (four T_0_ controls in total). To account for the proportion of the fuel that was biodegraded, we normalized hydrocarbon masses in ULSFO to the conserved internal marker 17α(H), 21β(H)-hopane (Prince et al., 1994). Given the low concentrations of hopane in diesel fuels, we used the nC17/pristane ratio to normalize the Marine diesel masses (Prince et al., 1994).

### Nucleic acid extraction

The netting preserved in DNA/RNA Shield was first thawed on ice prior to the nucleic acid extraction. The Falcon tubes were vortexed for 90 s to detach any bound cells and particles from the netting and resuspend them in the supernatant. The solutions were left to settle on ice until the foam produced by vortexing dissipated. DNA was then extracted using the ZymoBIOMICS DNA/RNA Miniprep Kit (Zymo Research) following the manufacturer’s instructions with minor modifications. The supernatant (250 µl) was added to the ZR BashingBead Lysis Tube and mixed with 750 µl DNA/RNA Shield and the lysis tubes were shaken for 5 min on a Mini-Beadbeater-16 (BioSpec Products). All centrifugation steps were carried out at 16,000 g unless stated otherwise by the original protocol and in the final elution step, DNA/RNA was eluted in 50 µl of ZymoBIOMICS DNase/RNase-Free Water. An extraction control consisting of 500 µl of ZymoBIOMICS DNase/RNase-Free Water and 500 µl of DNA/RNA Shield was processed along with the samples as described above. The resulting DNA extractions were stored at −20 °C until processed.

### Library preparation and sequencing

The 16S rRNA gene was amplified using primers 515F-Y (5′-GTGYCAGCMGCCGCGGTAA) and 926R (5′-CCGYCAATTYMTTTRAGTTT) containing Illumina overhang adapter sequences (Parada et al., 2016). PCR reactions (25 µL) containing 0.5 U of KAPA HiFi DNA Polymerase (Roche), 0.6 µM of each primer, 0.3 mM of KAPA dNTP Mix (Roche), 1X KAPA HiFi Fidelity Buffer (Roche), and 1 µL of DNA were performed under the following conditions: initial denaturation at 95°C for 5 min, followed by 25 cycles of 95°C for 30 s, 50°C for 30 s, 72°C for 45 s, and a final extension step at 72°C for 10 min. Reactions were purified using Sera-Mag Select magnetic beads (Cytiva) with a 0.8 bead-to-PCR volume ratio. Indexing was performed using the Nextera XT Index Kit v2 (Illumina) following manufacturer’s instructions. Indexed samples were purified with Sera-Mag Select magnetic beads (1.12 bead-to-PCR volume ratio) and quantified using the Qubit fluorometer (Invitrogen). Samples were pooled in equimolar ratios of 4 nM and sequenced with a 2 × 300 bp v3 flow cell with an Illumina MiSeq platform. Adapters and indices were removed with the Illumina FASTQ file generation pipeline.

### Bioinformatics and statistical analyses

All statistical analyses were performed in R 4.4.1(R Core Team, 2024).

#### Hydrocarbon analysis

We tested for differences in hydrocarbon removal between the fertilized and unfertilized samples using a paired t-test or a paired Wilcoxon signed-rank test when the assumption of normality was not met. We could not detect a statistical effect of the use of fertilizer on the amount of removed hydrocarbons (Fig. S1; Table S1). We also did not find statistical differences in the microbial community (see below). Accordingly, we combined the fertilized and unfertilized samples into their respective categories (e.g., Marine diesel intertidal, etc.) for the remainder of the analyses to increase the statistical power of the tests. We used a two-way ANOVA to test for differences between treatments and locations (intertidal vs. supratidal zones) followed by a Tukey’s honest significant differences post-hoc test.

### 16S rRNA gene amplicons

Amplicon sequence variants (ASVs) were obtained from the 16S rRNA gene reads using DADA2 v1.26.0 (Callahan et al., 2016a). ASVs were classified using the Silva 138.1 database (Quast et al., 2013). The negative control reads were used to remove potential contaminant ASVs with decontam v1.22.0 (Davis et al., 2018) using the frequency method. Mitochondrial, chloroplast, and ASVs unclassified at the phylum level were also removed. We averaged ASV abundances across 1000 rarefactions to the minimum library size of 5727 as described previously (Sugden et al., 2022). A phylogenetic tree was inferred using phangorn v2.11.1 (Schliep, 2011) using the method described elsewhere (Callahan et al., 2016b). The resulting ASV table was imported to phyloseq v1.46.0 (McMurdie and Holmes, 2013) for statistical analyses.

Shannon and Faith’s phylogenetic diversities were calculated for all samples. Faith’s phylogenetic diversity (Faith’s PD) was estimated using picante v1.8.2 (Kembel et al., 2010). We tested for differences in these two diversity metrics between nutrient amendments (fertilized vs. unfertilized) with a paired t-test or a paired Wilcoxon signed-rank test when the assumption of normality was not met. Similar to the hydrocarbon analyses, we did not observe any statistical changes in the microbial communities between the nutrient amendment treatments (Fig. S2; Table S2) so samples were combined as described above. Differences between treatments and locations were calculated with an ANOVA followed by Tukey’s honest significant differences post-hoc test. Bray-Curtis dissimilarities and weighted UniFrac distances were calculated to test for differences in community composition between treatments and locations with a PERMANOVA (Anderson, 2001) after using PERMDISP (Anderson and Walsh, 2013) to test for homogeneity in multivariate dispersions with vegan v2.6-4 (Oksanen et al., 2022). After obtaining significant results for the global PERMANOVA, we tested for pairwise differences among the categories of the significant variables using a pairwise PERMANOVA (Martinez Arbizu, 2020).We determined differentially abundant ASVs between treatments and locations with ANCOM-BC2 (Lin and Peddada, 2024). We evaluated the influence of aliphatic and polycyclic aromatic hydrocarbons on the microbial community with a distance-based redundancy analysis (dbRDA) in vegan.

## Results

### Hydrocarbon analysis of the residual fuel

After one year, we observed a significantly higher (ANOVA, F = 163.26, p < 2 × 10^-16^) removal (79.8 ± 4.1%) of the added Marine diesel compared to the 62.1 ± 20.3% of the ULSFO removed from the mesocosms (Fig. 1). There was a higher removal in the supratidal zone compared to the intertidal zone (ANOVA, F = 93.29, p = 3.13 × 10^-12^) for both Marine diesel and ULSFO. Tukey HSD test results showing differences among individual variable combinations presented below can be found in Table S3. After normalizing to account for biodegradation, we determined that 72.0 ± 15.8% of the Marine diesel was removed by biological processes and there were no statistical differences between the proportion of fuel removed by natural attenuation and by biodegradation for this fuel. On the other hand, only 32.5 ± 9.6% of the ULSFO was biodegraded and this value was significantly lower than the ULSFO removal by natural attenuation. The removal of Marine diesel was consistently higher than for ULSFO by both natural attenuation and biodegradation for the two tidal zones. The one exception was natural attenuation in the supratidal zone were there were no statistical differences between the fuels. Aliphatic hydrocarbon removal was higher in Marine diesel compared to the PAH removal. For ULSFO, there were no statistical differences between the removal of the two classes of compounds except for the natural attenuation in the intertidal zone for which there was a higher removal of aliphatics compared to PAHs.

**Fig. 1.**
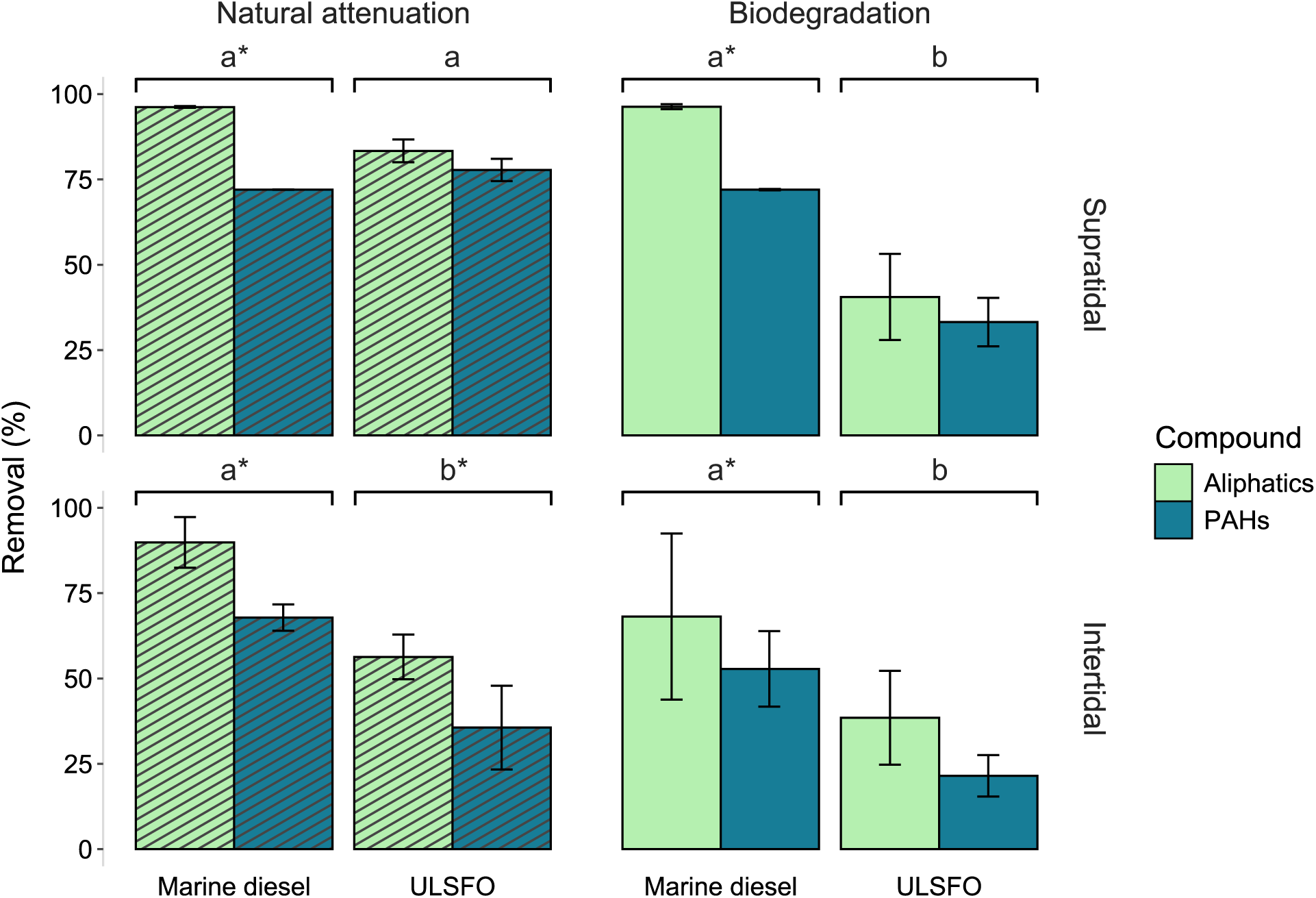
Removal percentages of the studied fuels by natural attenuation and biodegradation and by compound group. Letters represent statistical differences among the fuels for each of the panels. An asterisk (*) represent statistical differences in the removal percentage by compound group inside a given fuel and removal type combination.

### Changes in the beach microbiome

The phyla *Pseudomonadota* and *Bacteroidota* comprised the majority of the 2512 ASVs detected in the mesocosms (Fig. 2). The top 20 most abundant ASVs also belonged to these two phyla with the exception of ASV18 which belonged to the phylum *Verrucomicrobiota* (Fig. S3; Table S4). The ANCOM-BC2 results showed that bacteria from the classes *Bacteroidia* (phylum *Bacteroidota*), *Alphaproteobacteria* and *Gammaproteobacteria* (phylum *Pseudomonadota*), and *Verrucomicrobiia* (phylum *Verrucomicrobiota*) were more abundant in the treated samples compared to the untreated and T_0_ controls and the majority of these were observed in the supratidal zone (Fig. 3). When comparing whether there were any differentially abundant classes between the tidal zones of the same treatment, only the class *Alphaproteobacteria* was more abundant in the supratidal zone of the ULSFO treatment compared to the samples of this treatment in the intertidal zone (Fig. 3b). At the genus level, ANCOM-BC2 showed that 14 genera from the phyla *Pseudomonadota* and *Bacteroidota* were more abundant in the fuel treatments compared to the controls and 4 *Pseudomonadota* genera were more abundant in the controls compared to the treated samples (Fig. S4; Table S5). We also observed 13 genera from the phyla *Bacteroidota* and *Pseudomonadota* with higher abundances in the supratidal zone and 6 genera from the same two phyla that were more abundant in the intertidal zone (Fig. S5; Table S6).

**Fig. 2.**
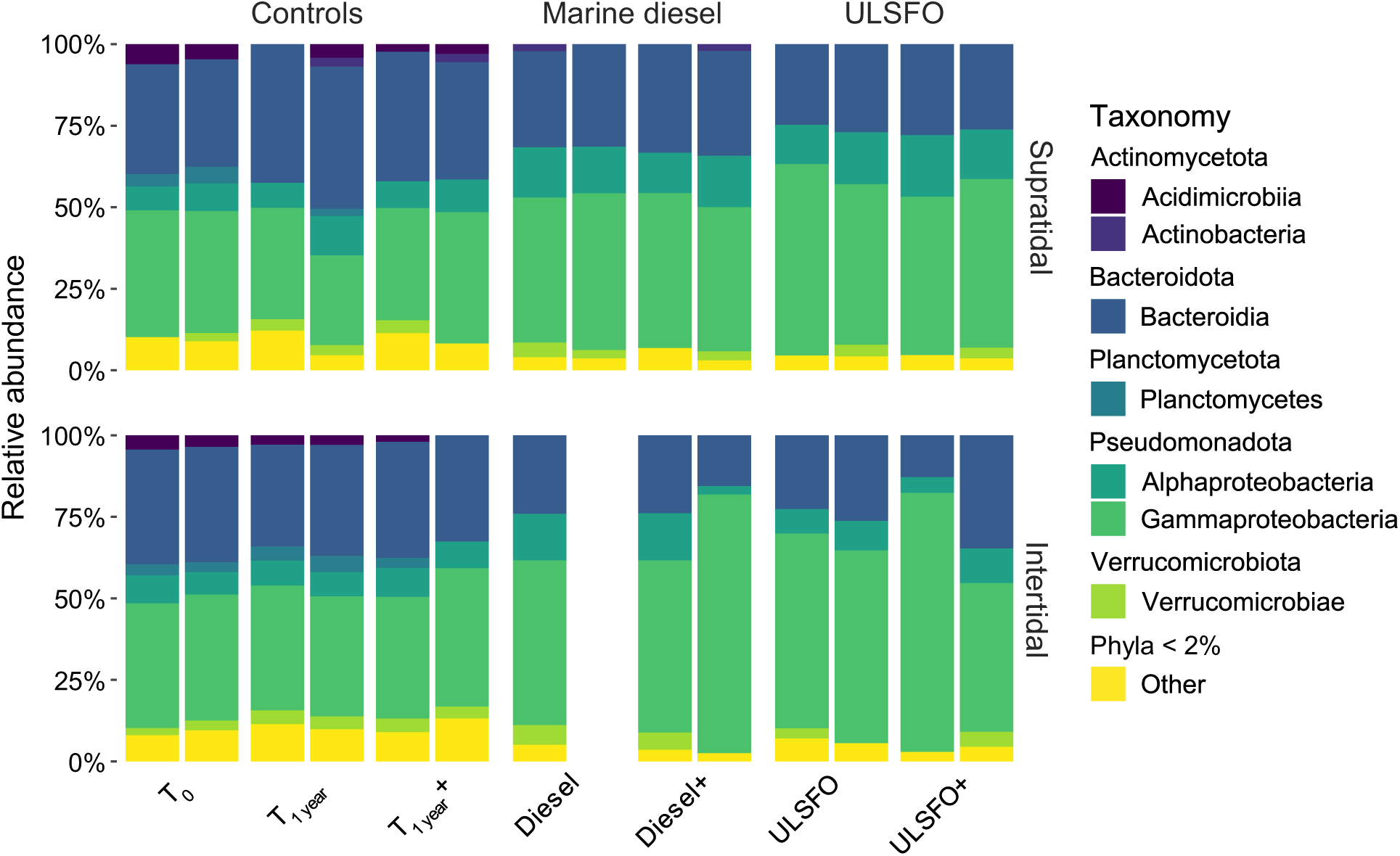
Taxonomy of the microbial communities of the *in situ* mesocosms after a year based on 16S rRNA gene amplicon sequencing. Phyla with a relative abundance lower than 2% were pooled into “Other”.

**Fig. 3.**
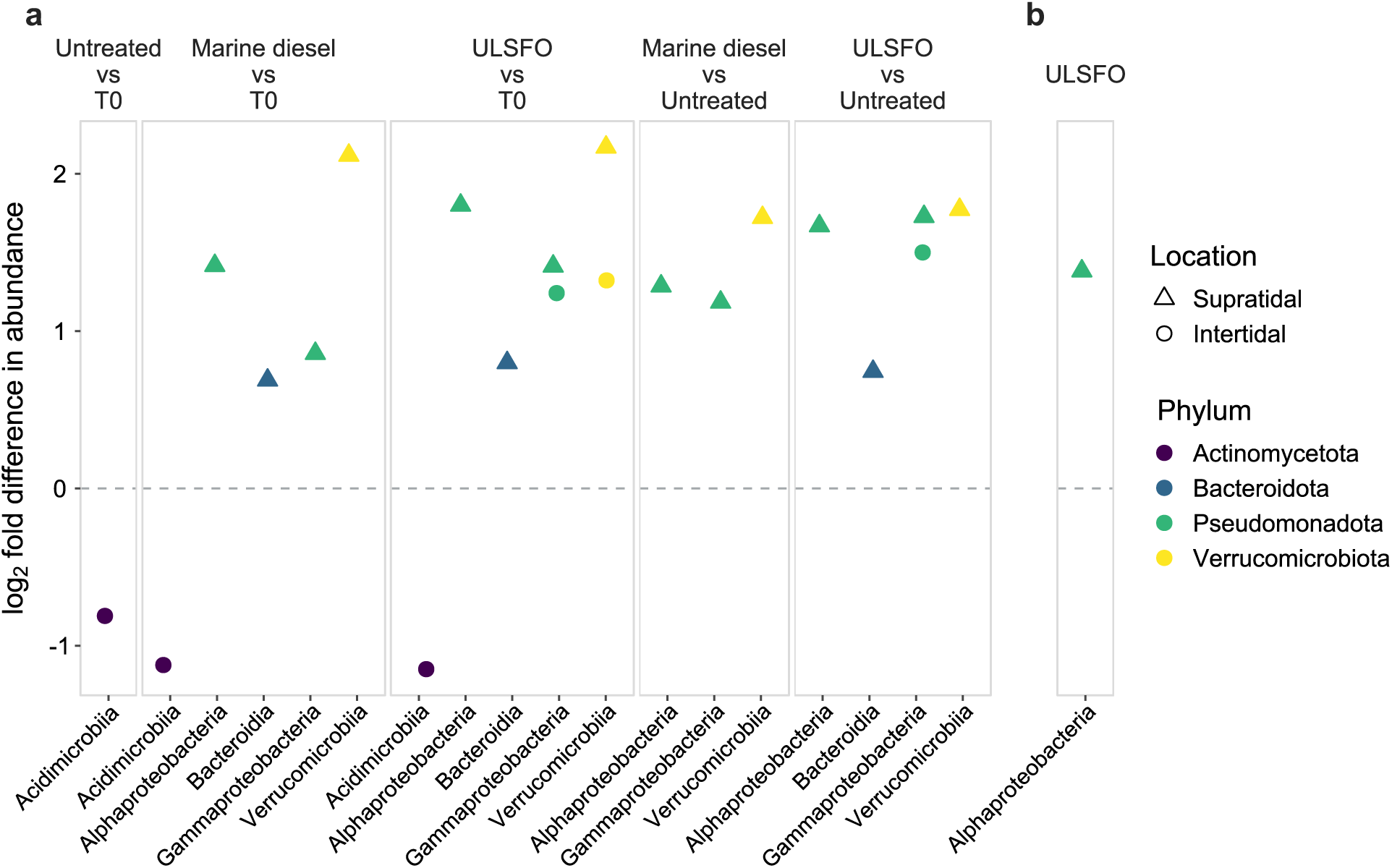
ANCOM-BC2 results showing the differentially abundant ASVs pooled by class. (**a**) Positive values represent ASVs that were more abundant in the first treatment in the contrast and negative values represent ASVs that were more abundant in the second treatment of the contrast. (**b**) Positive values represent ASVs that were more abundant in the supratidal zone and negative values represent ASVs that were more abundant in intertidal zone.

We observed a statistically significant decrease in both Shannon diversity and Faith’s PD for the samples treated with fuel compared to the controls for the supratidal zone and a significant decrease in Shannon diversity, but not in Faith’s PD for the intertidal zone (Fig. 4; Table S7). There were no statistical differences between the intertidal and supratidal samples within the same treatment. Similarly, we observed differences in community composition based on both Bray-Curtis dissimilarities and weighted UniFrac distances between the untreated controls and the fuel samples (Fig. 5; Table S8). However, unlike for the alpha diversity metrics, we did detect differences between the community composition between the intertidal and supratidal samples of the same treatment. The only pairwise comparison for which we did not obtain a statistical difference was for the weighted UniFrac distance between Marine diesel and ULSFO.

**Fig. 4.**
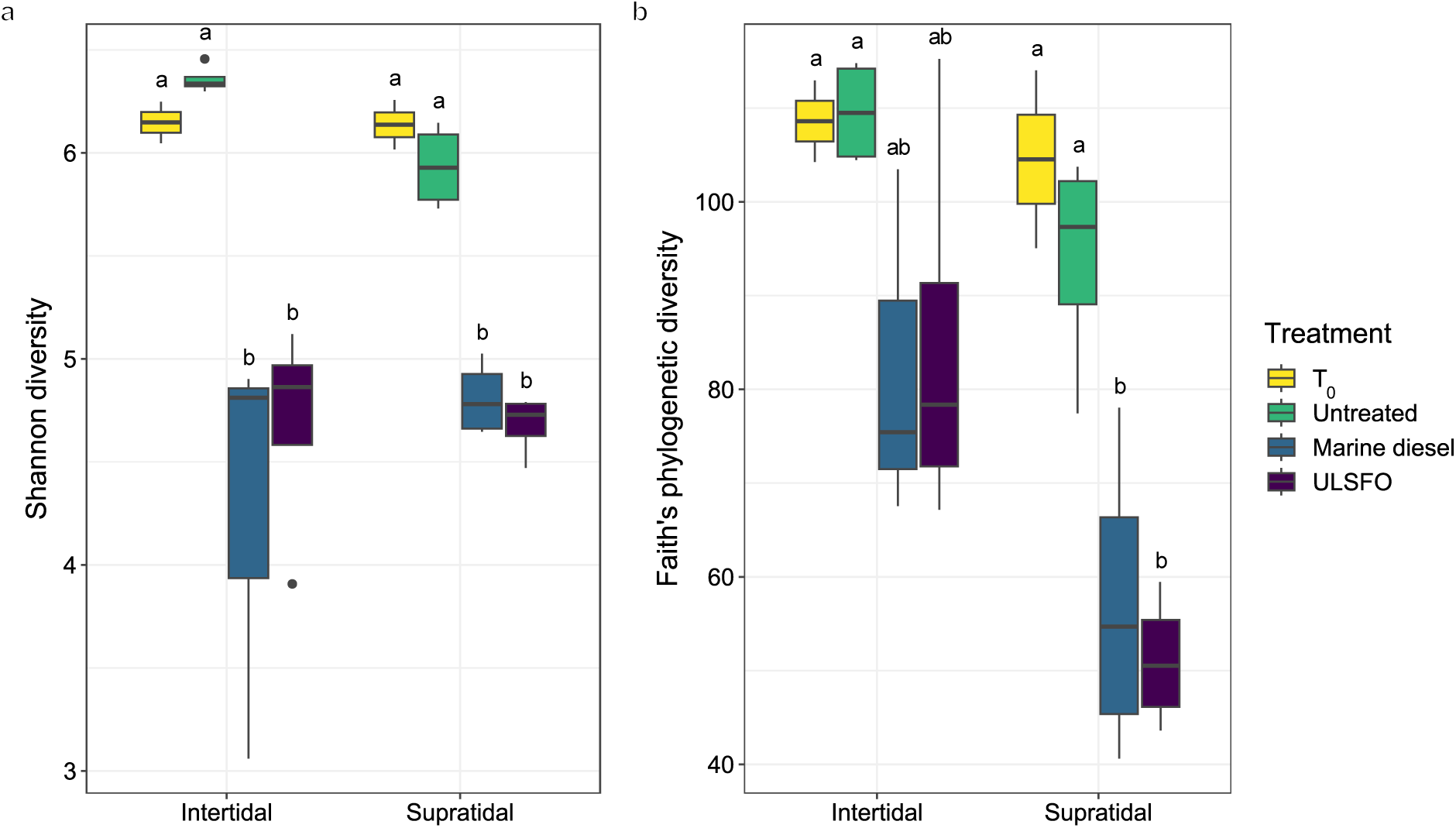
Differences in (**a**) Shannon diversity and (**b**) Faith’s phylogenetic diversity among the 16S rRNA gene amplicon community of the treatments.

**Fig. 5.**
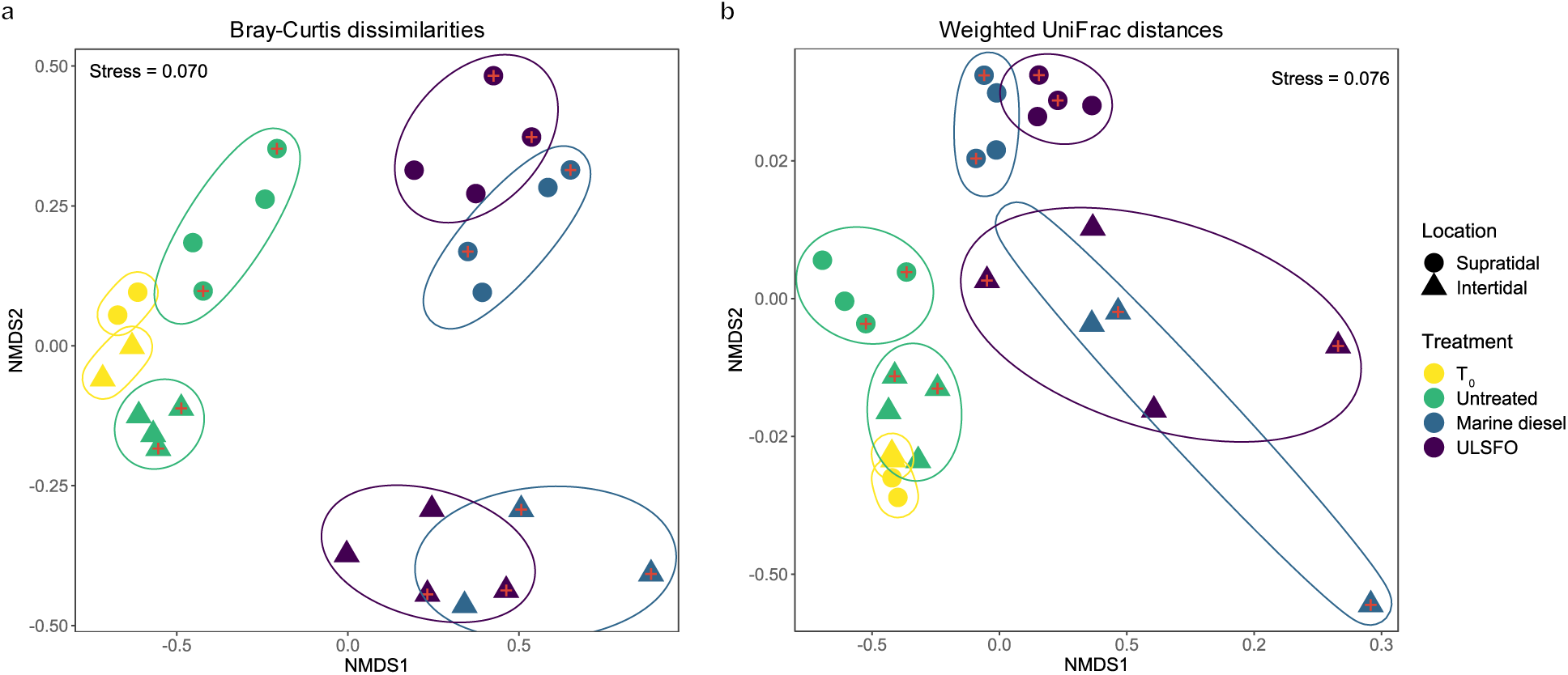
Non-metric multidimensional scaling (NMDS) ordination of Bray-Curtis dissimilarities (**a**) and (**b**) weighted UniFrac distances for the 16S rRNA gene amplicon community composition showing how communities adapted in response to the presence of the added fuels. Symbols with red crosses represent samples for which fertilizers were added at the beginning of the experiment.

The dbRDA further illustrated how the hydrocarbon concentrations explain the differences we observed in the microbial communities (Fig. 6). For the Bray-Curtis dissimilarities, the aliphatic concentrations had a significant effect on the community composition and the ULSFO samples were more strongly driven by these values, compared to the samples with Marine diesel (Fig. 6a). Both the aliphatic and PAH concentrations significantly influenced the community composition based on the weighted UniFrac distances (Fig. 6b). Similar to the Bray-Curtis dbRDA, the ULSFO microbial communities were more strongly driven by the hydrocarbon concentrations compared to the Marine diesel samples. Additionally, the dbRDAs illustrated that the community composition of the T_0_ and untreated controls were negatively correlated to the aliphatic and PAH concentrations.

**Fig. 6.**
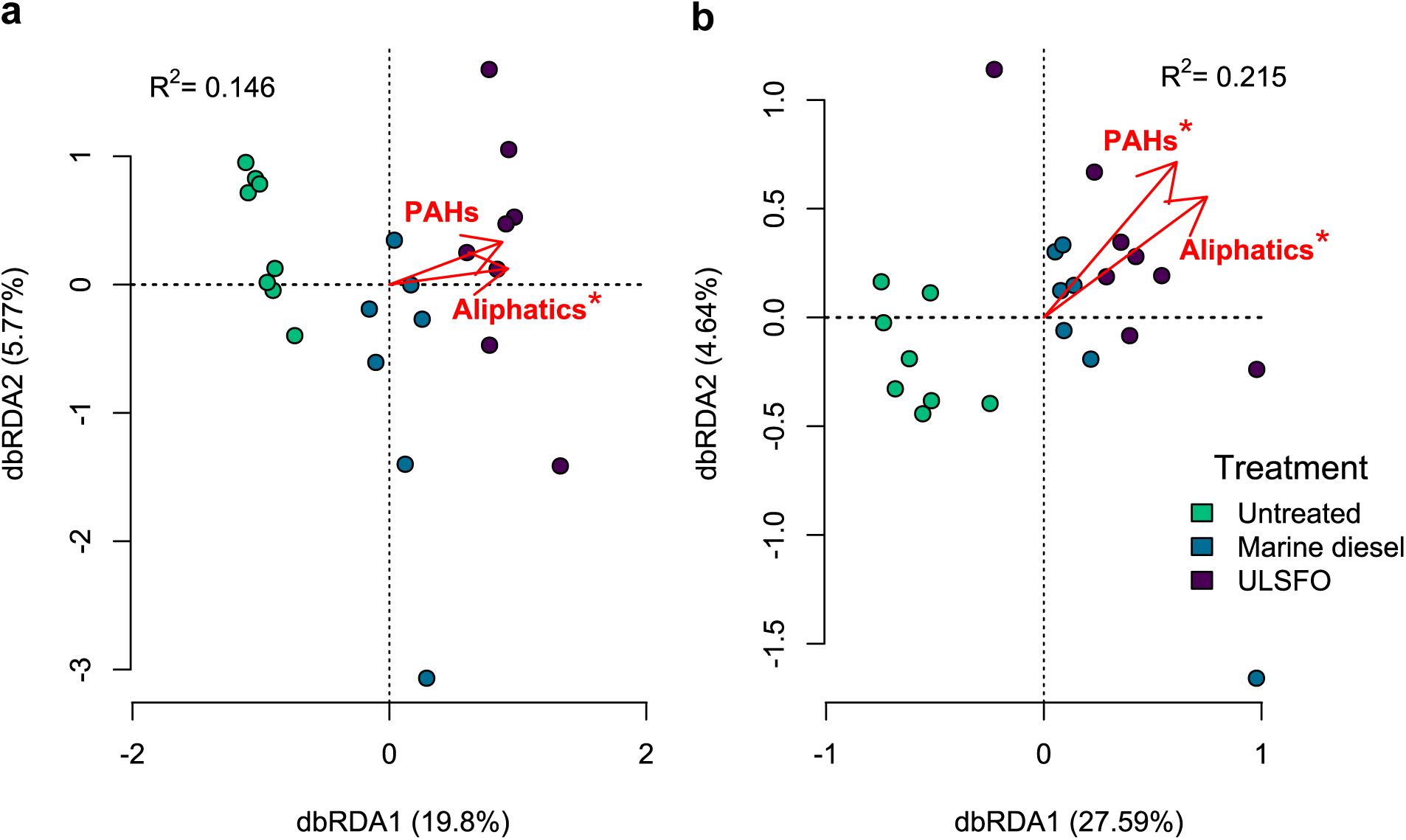
Distance-based redundancy analysis (db-RDA) illustrating the influence of the type of hydrocarbon on the 16S rRNA gene amplicons for (**a**) Bray-Curtis dissimilarities- and (**b**) weighted UniFrac-based community composition.

## Discussion

Previous work at Assistance Bay (Góngora et al., 2024a) showed that the microbial community of this NWP beach was capable of degrading only a portion of the applied fuel after 33 days. In the present study, we followed up the one-month experiment by deploying the *in situ* mesocosms for a full year to determine whether the extended contact time along with the addition of nutrient amendments improved hydrocarbon removal.

We did not observe an effect of the use of fertilizers in hydrocarbon removal (Fig. S1), in the alpha diversity metrics (Fig. S2) or in the community composition (Fig. 5). Previous hydrocarbon biodegradation studies using fertilizer amendments in high Arctic beach sediments obtained mixed results. Ellis et al. (2022) observed an increase in hexadecane radiorespiration rates in nutrient amended microcosms compared to the unamended samples only for one of the three studied beaches and the addition of fertilizers did not improve the naphthalene radiorespiration rates. The authors also saw no differences in ULSFO removal between the fertilized and unfertilized microcosms. In a laboratory experiment simulating tide activity with sediment from a beach in Resolute, Y.-J. Chen et al. (2024) reported an increase in alkane removal after 32 days for the treatment containing ULSFO, MAP, and S200 compared to the fuel only treatment, but did not detect an effect of the nutrient amendments after 92 days. The *In situ* Treatment of Oiled Sediment Shorelines program carried out in a shoreline in Svalbard, Norway (1997–1998) estimated that fertilizer treatments applied during the first two months of the experiment doubled biodegradation rates of the intermediate fuel oil IF-30 after a year (Prince et al., 2003). The Baffin Island Oil Spill (BIOS) project carried out in Baffin Island, Canada (1980-1984) saw an improvement in biodegradative activity of Lago Medio crude oil for the treatment plots supplemented with fertilizers (Eimhjellen and Josefsen, 1984). One explanation for the lack of effect of fertilizers in our study could be the fact that we only added these nutrient amendments once at the beginning of the experiment. It is then possible that the fertilizer was consumed very quickly by the microbial community in the first days of the experiment and we could not observe the short-term effect with the sampling points selected. This is consistent with the experiment by Y.-J. Chen et al. (2024) which saw a biostimulation effect only in the first time point or with the BIOS project which determined that nutrient amendments increased the biodegradation rates, but not the end-point oil loadings (Owens et al., 2003). Another possibility is that the added fertilizers in our experiment were washed away by tidal action within the first couple of tidal cycles. This is why it has been suggested that constantly monitoring nutrient and adding fertilizers more than once could lead to an optimal biostimulation treatment (Prince et al., 2003).

After a year, we observed a statistically higher hydrocarbon removal of both fuels in the supratidal zone compared to the intertidal zone (Table S3). For Marine diesel, the lack of statistical differences between the natural attenuation and biodegradation percentages suggests that most of the removal (72.0%) can be attributed to the metabolic activity of hydrocarbon degraders. On the other hand, we did observe statistical differences between the natural attenuation and biodegradation (32.5%) removal of ULSFO meaning that a substantial proportion (29.6%) of this fuel was removed by non-biological processes. Some of the differences that we observed could be explained by the type of hydrocarbons present in the used fuels (Fig. S6). Marine diesel is mostly comprised of aliphatic hydrocarbons (93.3%) while 26.4% of the quantified compounds in ULSFO are PAHs. Our previous studies of the microbial community of Assistance Bay found that there is limited abundance and expression of genes associated with PAH degradation on the microbiome of this beach (Freyria et al., 2024; Góngora et al., 2024a, 2024b) as well as in other beaches across the NWP (Durand et al., 2023; Ellis et al., 2022).

We observed a pronounced difference in the microbial communities of the fuel-treated samples compared to the T_0_ and untreated controls (Fig. 5; Table S8). This separation between the treated and untreated samples evidences the capacity of the Assistance Bay beach sediment microbial community to adapt to the sudden addition of hydrocarbons as was also observed with the dbRDA which showed a strong correlation of the treated samples with the hydrocarbon concentrations (Fig. 6). The ANCOM results also showed that four of the 20 most abundant genera (*Oleispira*, *Altererythrobacter*, *Gilvibacter*, *Pseudohongiella*) were differentially more abundant in the fuel treated samples compared to the untreated controls (Table S5). However, the differences also appear to be caused by a reduction in diversity (Fig. 4; Table S7) showing that the added fuels could have also negatively affected the baseline microbiome while enriching for hydrocarbon degraders. These statistically significant differences in alpha diversity and community composition were not observed in the one-month experiment (Góngora et al., 2024a) which could suggest that the environmental impacts of a fuel spill in the NWP are greater for the microbiota after a longer exposure.

It is also worth noting that there were no differences in Faith’s PD between the fuel treatments and the controls for the intertidal zone, but there were differences in weighted UniFrac for these samples. These results could be indicating that the microbial community in the intertidal zone is changing very slowly. The weighted UniFrac NMDS also illustrate how there is a larger dispersion in the fuel-treated intertidal samples compared to their supratidal counterparts (Fig. 5B) which further supports that the microbial communities in these samples have not fully stabilized. Additionally, there is a larger number of differentially abundant genera in the supratidal zone compared to the intertidal zone (Fig. S5; Table S6) which indicates that bacteria from the same genus appear to thrive better in the supratidal zone. We can see that 38.5% of the genera that were most abundant in the supratidal zone are known hydrocarbon degraders, while only 16.7% of the genera that were more abundant in the intertidal zone were hydrocarbon degraders (Table S6). On the other hand, 66.7% of the genera more abundant in the intertidal zone (23.1% in the supratidal zone) have been associated with hydrocarbon degradation, but their biodegradative capabilities have never been confirmed. There have been various documented cases suggesting that the presence of a given taxa inside the microbial community of a hydrocarbon spill does not necessarily mean that said taxa is capable of hydrocarbon biodegradation. For example, *Colwellia* which was differentially more abundant in the supratidal zone of the ULSFO treatments, does not appear to possess or express genes involved in hydrocarbon degradation based on results of microcosm experiments using water from the water plume of the Deepwater Horizon oil spill (Peña-Montenegro et al., 2023). The transcriptomic activity of bacteria from this genus led the authors to hypothesize that *Colwellia* might be opportunistic bacteria, rather than a primary hydrocarbon degrader. Hydrocarbon degradation genes were scarcely distributed across various *Lutibacter* genomes, a genus that was more abundant in the intertidal zone of the T_0_, Marine diesel, and ULSFO mesocosms. The authors suggested that this and other genera with few hydrocarbon activation genes could be secondary consumers that use the intermediary metabolic products produced by primary hydrocarbon degraders (S.-C. Chen et al., 2024). A limited colonization of the intertidal mesocosms by hydrocarbon degraders could thus explain why there was less biodegradation compared to the supratidal zone (Fig. 1; Table S3).

While our results provide some of the first evidence of the *in situ* biodegradation of ULSFO under high Arctic environmental conditions, not much is known about the behaviour and properties of LSFOs in general. Each LSFO has a different chromatographic signature (Daling and Sørheim, 2020; Faksness et al., 2024; Nelson et al., 2022; Yang et al., 2023) which could be beneficial for tracing and identification of a spill from an unknown fuel source, but it could also present issues in determining whether biodegradation could be a feasible remediation solution since the microbial communities could respond differently to each type of LSFO. For example, two different batches of ULSFO produced by Shell contain different proportions of long-chain alkanes (Daling and Sørheim, 2020). LSFOs have been seen to become very viscous at low temperatures and viscosity further increases when water-oil emulsions are formed (Lee et al., 2023). The Swedish Coast Guard reported that after a VLSFO spill off the coast of Sweden, the fuel tended to cluster into small lumps when it reached the shore (Pålsson et al., 2024). ULSFOs have a high (> 12%) wax content (Daling and Sørheim, 2020; IMAROS, 2022) which further increases the viscosity of these fuels. While the increased viscosity could reduce fuel penetration into the beach sediment, it may also hamper biodegradation as it reduces the surface area where microorganisms come into contact with the hydrocarbons.

## Conclusions

The limited amount of research on LSFOs presents a high risk for governments and response teams across the world as there does not appear to be a reliable cleanup strategy in case a LSFO spill washes onto a shoreline. In this study, we aimed to help to close the knowledge gap by evaluating whether nutrient biostimulation could improve the natural attenuation removal capacity of the microorganisms inhabiting a high Arctic beach in the NWP. We observed that there is indeed a natural biodegradative capacity in the Assistance Bay shoreline microbiome, even if we did not achieve a complete removal of the applied fuel. On the other hand, we observed higher biodegradation for Marine diesel which could be caused by its less complex, alkane-rich nature. We also observed that a single fertilizer application at the beginning of the year-long experiment was not sufficient to provide a long-term improvement in performance to the already present natural attenuation potential of the microbes inhabiting the shoreline. These findings show that the native beach microbiome does not appear to be capable of fully degrading a ULSFO spill if one were to happen on a NWP shoreline. A higher level of monitoring to help determine if more frequent fertilization application events might be needed along with more invasive stimulation techniques such as the application of surface washing agents or mechanical removal are required to achieve a complete cleanup response.

## CRediT authorship contribution statement

**Esteban Góngora:** Conceptualization, Data curation, Formal analysis, Funding acquisition, Investigation, Methodology, Visualization, Writing – original draft, Writing – review and editing. **Ya-Jou Chen:** Conceptualization, Investigation, Methodology, Writing – review and editing.

**Nastasia J. Freyria:** Investigation, Methodology, Writing – review and editing.

**Antoine-O. Lirette:** Investigation, Methodology, Writing – review and editing.

**Charles W. Greer:** Funding acquisition, Supervision, Writing – review and editing.

**Lyle G. Whyte:** Funding acquisition, Project administration, Resources, Supervision, Writing – review and editing.

## Acknowledgements

This work was funded by Fisheries and Oceans Canada and Natural Resources Canada under the Oceans Protection Plan’s Multi-Partner Research Initiatives and by the Fonds de recherche du Québec - Nature et technologies (No. 273122). Arctic logistical support was provided by the Polar Continental Shelf Program from Natural Resources Canada and the Northern Scientific Training Program from Polar Knowledge Canada. We would like to thank Devon Manik from Resolute who was our guide and bear watcher during the deployment of the mesocosms.

