## Supplemental figures for "Long-term patterns of ultra-low sulfur fuel oil bioremediation in Arctic shorelines using *in situ* mesocosms"

[†] Current affiliation: Department of Environmental Microbiology, Institute for Sanitary Engineering, Water Quality and Solid Waste Management (ISWA), University of Stuttgart, Stuttgart, Germany

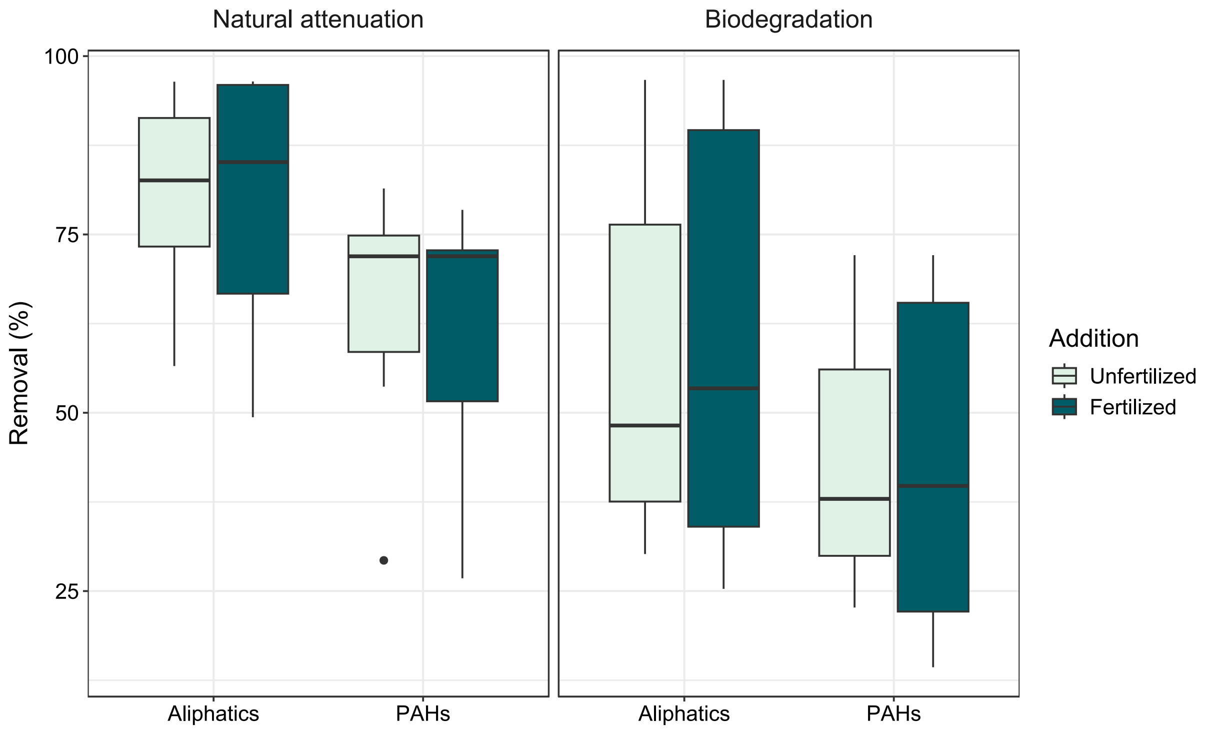


**Fig. S1.** Removal percentages of the studied fuels by natural attenuation and biodegradation for the fertilized and unfertilized samples.

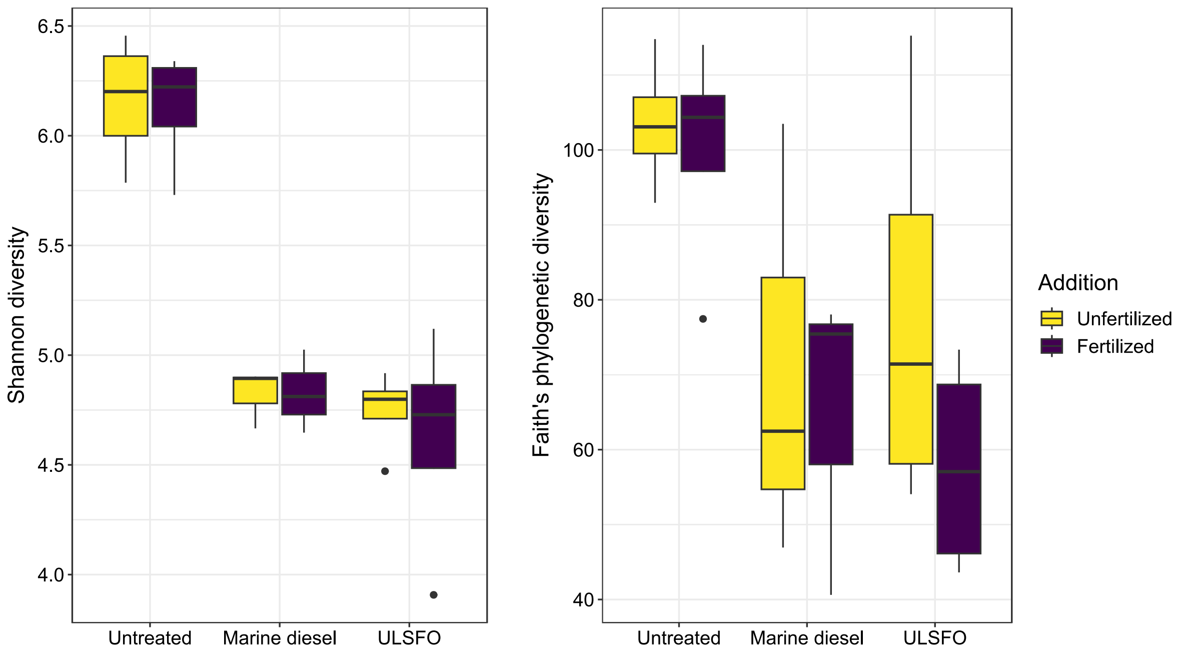


**Fig. S2.** Differences in Shannon diversity and Faith's phylogenetic diversity between the 16S rRNA gene amplicon communities of the fertilized and unfertilized samples.

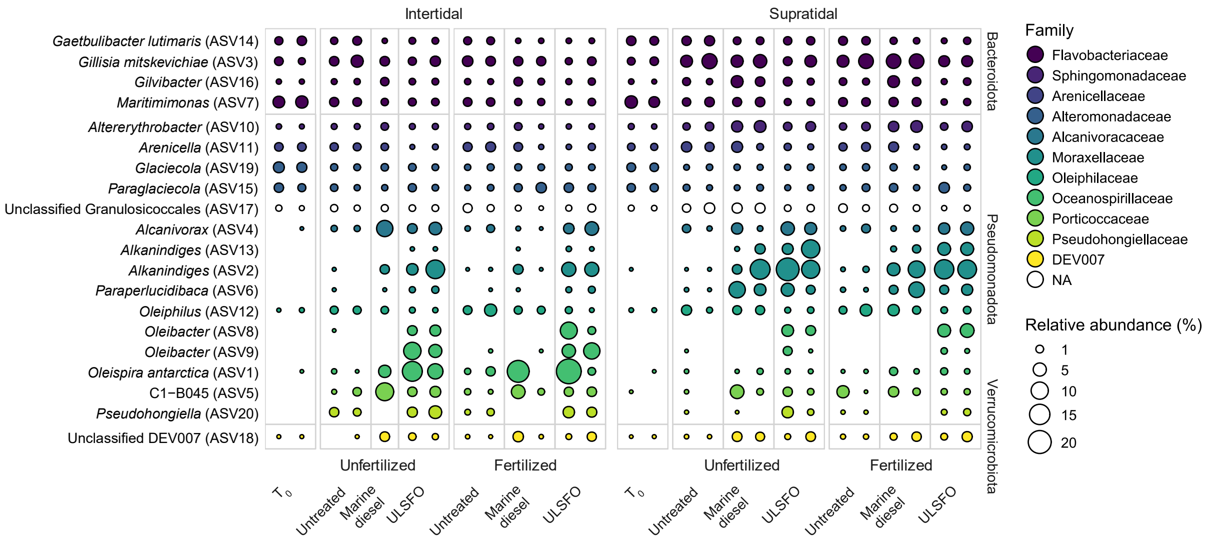


**Fig. S3.** Relative abundances of the top 20 most abundant ASVs detected based on 16S rRNA gene amplicon community.

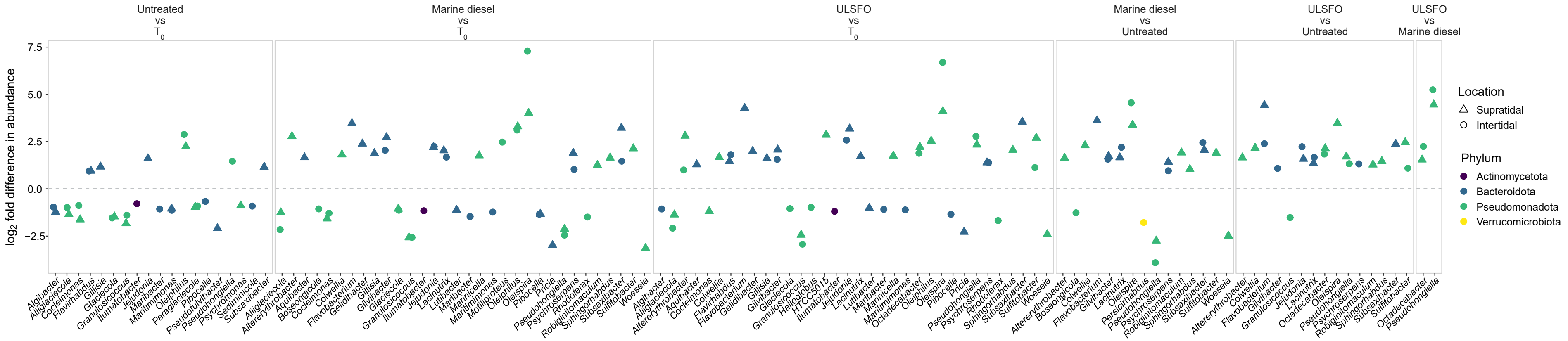


**Fig. S4.** ANCOM-BC2 results showing the differentially abundant ASVs detected among treatments by genus. Positive values represent ASVs that were more abundant in the first treatment in the contrast and negative values represent ASVs that were more abundant in the second treatment of the contrast.

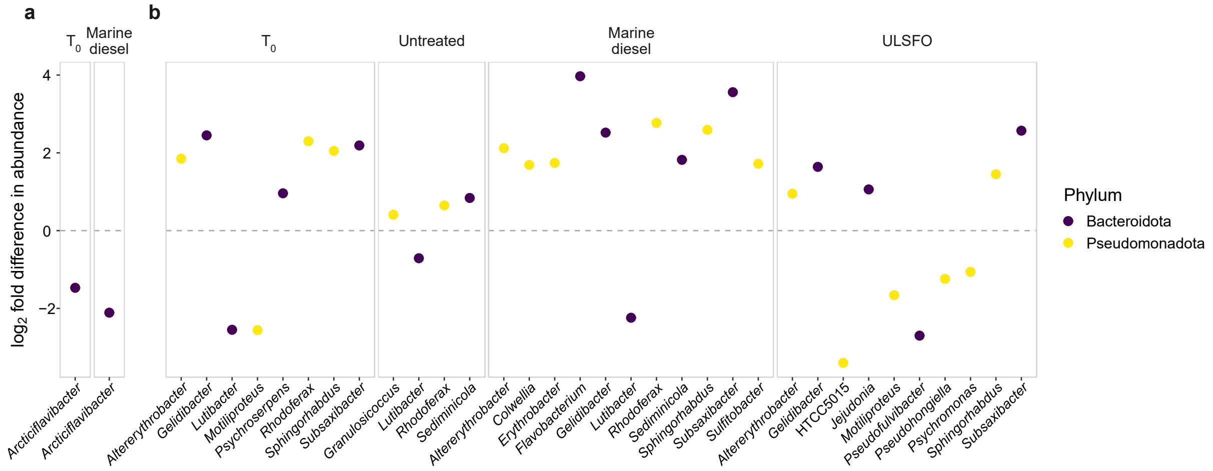


**Fig. S5.** ANCOM-BC2 results showing the differentially abundant ASVs pooled by genus detected (**a**) between fertilized and unfertilized samples and (**b**) between the intertidal and supratidal communities. For (**a**) positive values represent ASVs that were more abundant in the fertilized samples and negative values represent ASVs that were more abundant in the unfertilized samples. For (**b**) positive values represent ASVs that were more abundant in the supratidal zone and negative values represent ASVs that were more abundant in the intertidal zone.

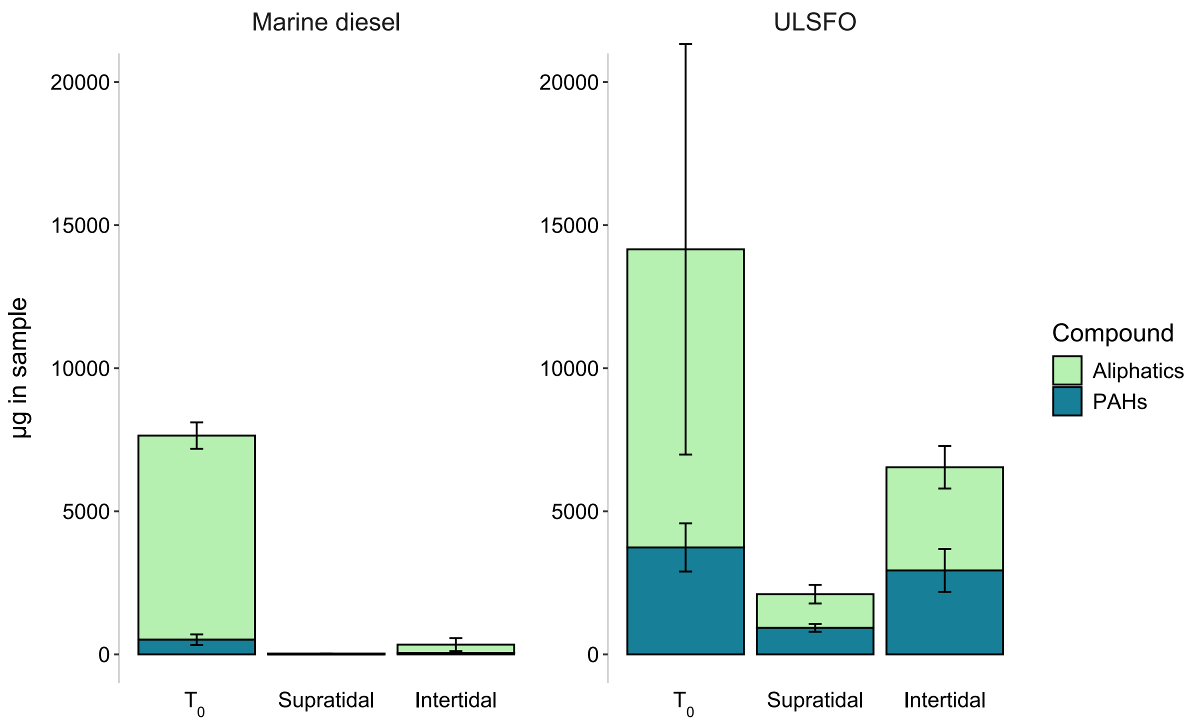


**Fig. S6.** Initial and final hydrocarbon masses of the studied fuels by type of compound.
